# Novel synergies and isolate specificities in the drug interactions landscape of *Mycobacterium abscessus*

**DOI:** 10.1101/2022.12.12.520102

**Authors:** Nhi Van, Yonatan N. Degefu, Pathricia A. Leus, Jonah Larkins-Ford, Jacob Klickstein, Florian P. Maurer, David Stone, Husain Poonawala, Cheleste M. Thorpe, Trever C. Smith, Bree B. Aldridge

## Abstract

*Mycobacterium abscessus* infections are difficult to treat and are often considered untreatable without tissue resection. Due to the intrinsic drug-resistant nature of the bacteria, combination therapy of three or more antibiotics is recommended. A major challenge in treating *M. abscessus* infections is the absence of a universal combination therapy with satisfying clinical success rates, leaving clinicians to treat infections using antibiotic lacking efficacy data. We systematically measured drug combinations in *M. abscessus* to establish a resource of drug interaction data and identify patterns of synergy to help design optimized combination therapies. We measured approximately 230 pairwise drug interactions among 22 antibiotics and identified 71 synergistic pairs, 54 antagonistic pairs, and four potentiator-antibiotics not previously reported. We found that commonly used drug combinations in the clinic, such as azithromycin and amikacin, are antagonistic in lab reference strain ATCC19977, whereas novel combinations, such as azithromycin and rifampicin, are synergistic. Another challenge in developing universally effective multidrug therapies for *M. abscessus* is the significant variation in drug response between isolates. We measured drug interactions in a focused set of 36 drug pairs across a small panel of clinical isolates with rough and smooth morphotypes. We observed highly strain-dependent drug interactions that cannot be predicted from single-drug susceptibility profiles or known drug mechanisms of action. Our study demonstrates the immense potential to identify synergistic drug combinations in the vast drug combination space and emphasizes the importance of strain-specific combination measurements for designing improved therapeutic interventions.

## INTRODUCTION

*Mycobacterium abscessus (M. abscessus)* is a rapidly growing nontuberculous mycobacterium (NTM) that consists of three subspecies, *M. abscessus* subsp. *abscessus*, *M. abscessus* subsp. *massiliense* and *M. abscessus* subsp. *bolletii* (1). *M. abscessus* is a ubiquitous, opportunistic pathogen commonly found in soil, water systems, and contaminated material in hospitals. Recently, a steady increase in the morbidity and mortality of NTM infections has been reported worldwide (2). Although *M. abscessus* causes both pulmonary and extrapulmonary infection, the majority of the clinical syndrome of *M. abscessus* infections are pulmonary, occurring in immunocompromised people and those with pre-existing conditions such as cystic fibrosis, chronic obstructive pulmonary diseases, and bronchiectasis (3, 4).

Pulmonary *M. abscessus* infections are notoriously hard to treat due to the plethora of intrinsic mechanisms conferring resistance toward most clinically relevant antimicrobials, including macrolides, aminoglycosides, tetracyclines, and ß-lactams (5). Antibiotics such as amikacin and moxifloxacin are bactericidal in *E. coli* but are bacteriostatic in *M. abscessus*, creating another barrier to treating *M. abscessus* infection (6). The inherent phenotypic drug resistance in *M. abscessus* is due to multiple factors. *M. abscessus* possesses complex drug efflux pump system. a cell wall rich in lipids and mycolic acids that act as a physical barrier between antibiotics and the bacteria (7). Additionally, *M. abscessus* express numerous enzymes that can modify drugs or their targets. For example, studies have identified multiple enzymes in *M. abscessus* such as the Erm(41) erythromycin ribosomal methyltransferase, aminoglycoside acetyltransferases, an aminoglycoside phosphotransferase, a rifamycin ADP-ribosyltransferase (Arr_Mab_), a ß-lactamase (Bla_mab_), and tetracycline-modifying monooxygenase that confer their ability to resist many clinically used antibiotics (7–9). In addition to intrinsic resistance, acquired mutational resistance against macrolides and aminoglycosides also poses a significant risk for chronically infected patients with increased long-term antibiotic exposure (10). These characteristics contribute to the complex and multifaceted resistome of *M. abscessus*, creating a major challenge in developing antimicrobial therapies against *M. abscessus*.

Despite being an emerging global health threat, pan-effective treatment for NTM lung diseases has not yet been established and current treatment guideline is based on small studies and expert opinion (11, 12). Hence, treatment of *M. abscessus* is highly individualized and typically split into an intensive initial phase of several weeks comprising at least three antibiotics and a continuation phase of several months. In patients infected with strains harboring inducible or mutational resistance to macrolides, a macrolide can be included in the regimen for its immunomodulatory purpose only (12). Current guidelines suggest a minimum treatment duration of twelve months after culture conversion, although it is also individualized based on several factors, such as underlying patient conditions and *M. abscessus* subspecies, which typically have different clinical outcomes (3, 13). Clinical outcomes are poor in patients with pulmonary disease and those with immunosuppression (2, 14). Meta-analysis done across multiple clinical studies showed that sputum conversion without relapse was low, and evidence suggests that surgical resection prolongs negative culture (13, 15, 16). The poor cure rate and the emergence of clinically acquired pan-macrolide and pan-aminoglycoside resistance suggest the urgent need to develop more effective therapies for NTM and *M. abscessus* infections (8). Except for clarithromycin, results of traditional antimicrobial susceptibility testing by the Clinical and Laboratory Standards Institute (CLSI) are not linked to clinical outcomes (17). Additionally, there are no published breakpoints for tigecycline, or repurposed MDR-TB drugs like clofazimine or bedaquiline that are used to treat *M. abscessus*. Because *M. abscessus* treatment requires multi-drug therapy, systematic interrogation of drug combinations and drug-drug interactions has the potential to identify synergistic combinations that will form the basis for more effective therapies.

Several combination studies for *M. abscessus* have been reported using traditional checkerboard assays. However, conducting large-scale drug combination measurements is not practical because systematic measurements of drug-dose combinations in the checkerboard make these assays too resource-intensive. Here, we utilized DiaMOND (diagonal measurement of n-way drug interaction), a measurement and analysis pipeline based on an efficient geometric sampling of a traditional checkerboard assay, to significantly decrease the number of measurements required in checkerboard assays (18). We modified the DiaMOND assay to measure potentiation effects of drugs that are not active on their own but may increase the efficacy of other compounds. Because there are few antibiotics that are active in *M. abscessus*, potentiator screens may be critical to developing combination therapies in *M. abscessus* and other NTMs. Using medium-throughput measurement, we generated a systematic, large-scale catalog of drug interactions for *M. abscessus* as a resource and evaluated drug interaction patterns. Our dataset includes antibiotics commonly used in the clinic and next-generation antibiotics, such as bedaquiline or benzimidazole SPR719, an active moiety of a non-fluoroquinolone gyrase inhibitor SPR720. We identified many novel synergistic drug pairs, suggesting the potential for effective multidrug regimens in the combination drug space. However, drug interactions were highly variable among different clinical isolates highlighting the importance of making isolate-specific drug combination measurement rather than searching for a universal drug combination to treat *M. abscessus* infection.

## RESULTS

### Design of systematic drug interaction study in ATCC19977

To generate a systematic dataset of drug interaction profiles comparable to other published drug interaction studies, we began by measuring drug combination responses in the *M. abscessus* reference strain ATCC19977, an *M. abscessus* subsp. *abscessus* variant showing inducible macrolide resistance due to a functional Erm(41) ribosomal methylase (19). We measured pairwise combination effects among 22 antibiotics drawn from four categories (Table 1): (**a**) antibiotics currently recommended for *M. abscessus* treatment such as amikacin, clarithromycin, and azithromycin, (**b**) anti-tuberculosis drugs such as linezolid and ethambutol, (**c**) drugs in development for *M. abscessus* infection such as bedaquiline and SPR719, and (**d**) drugs that show no significant effect on their own but can be used to potentiate other drugs, such as avibactam.

**Table 1.** Summary of antimycobacterial used in the study and their antimicrobial activity against *M. abscessus* ATCC19977.

|  | antimycobacterial | general mechanism of action | abbreviation | IC<sub>50</sub> (µg/mL) | IC<sub>90</sub> (µg/mL) |
| --- | --- | --- | --- | --- | --- |
| active drug | **amikacin** | protein synthesis | AMK | 6.6 | 110 |
|  | **amoxicillin** | cell wall | AMX | 300 | NA |
|  | **azithromycin** | protein synthesis | AZM | 4.3 | 680 |
|  | **bedaquiline** | respiration | BDQ | 0.16 | 1.3 |
|  | **cefoxitin** | cell wall | FOX | 9 | 28 |
|  | **cerulenin** | cell wall | CER | 2.2 | 7 |
|  | **clarithromycin** | protein synthesis | CLR | 0.45 | 80 |
|  | **ethambutol** | cell wall | EMB | 31 | 1200 |
|  | **levofloxacin** | DNA | LXF | 13 | 200 |
|  | **linezolid** | protein synthesis | LZD | 14 | 260 |
|  | **moxifloxacin** | DNA | MXF | 1.5 | 35 |
|  | **nitrofurantoin** | DNA | NFT | 44 | 1800 |
|  | **rifabutin** | RNA | RFB | 1.8 | 23 |
|  | **rifampicin** | RNA | RIF | 16 | 2100 |
|  | **SPR719** | DNA | SPR | 0.39 | NA |
|  | **telithromycin** | protein synthesis | TEL | 10 | NA |
|  | **thioridazine** | respiration | TZ | 23 | 39 |
|  | **vancomycin** | cell wall | VAN | 9 | 250 |
| potentiator | **avibactam** | β-lactamase inhibitor | AVI | NA | NA |
|  | **streptomycin** | RNA | STR | NA | NA |
|  | **tetracycline** | RNA | TET | NA | NA |
|  | **verapamil** | efflux pump inhibitor | VER | NA | NA |

Drug combinations may be constructed from antibiotics that are active on their own and are effective together. We may then search for synergistic drug combinations, e.g., sets of antibiotics that are more effective together than expected based on their single-antibiotic efficacies. Nevertheless, applying this approach systematically to *M. abscessus* poses a challenge due to its inherent drug resistance and the large-scale, resource-intensive, traditional checkerboard assays. To overcome the latter, we used DiaMOND to measure pairwise combination effects more efficiently. DiaMOND is an experimental analysis method approximating the checkerboard assay to measure drug interactions. The first step of DiaMOND is to determine the 50% and 90% growth inhibitory concentrations (IC_50_ and IC_90_, respectively) of every drug to design an equipotent combination dose-response curve. We sampled 20-25 doses to obtain a dose-response curve for every drug. From obtained IC_50_ values, we designed pairwise DiaMOND experiments for equipotent combinations of drug pairs (diagonal in Fig. 1A). Fractional inhibitory concentrations at 50% growth inhibition (FIC_50_) were calculated based on Loewe Additivity and Bliss Independence null models. Metrics from both models were strongly correlated (R^2^ = 0.75, Pearson correlation), so we focused on FICs calculated by Loewe Additivity (Fig. S1). We report log_2_FIC values, such that synergies (negative) are balanced in magnitude compared to antagonisms (positive). We can also visualize synergies and antagonisms by comparing combination dose-response curves to single drug dose-response curves (Fig. 1C)

**FIG 1.**
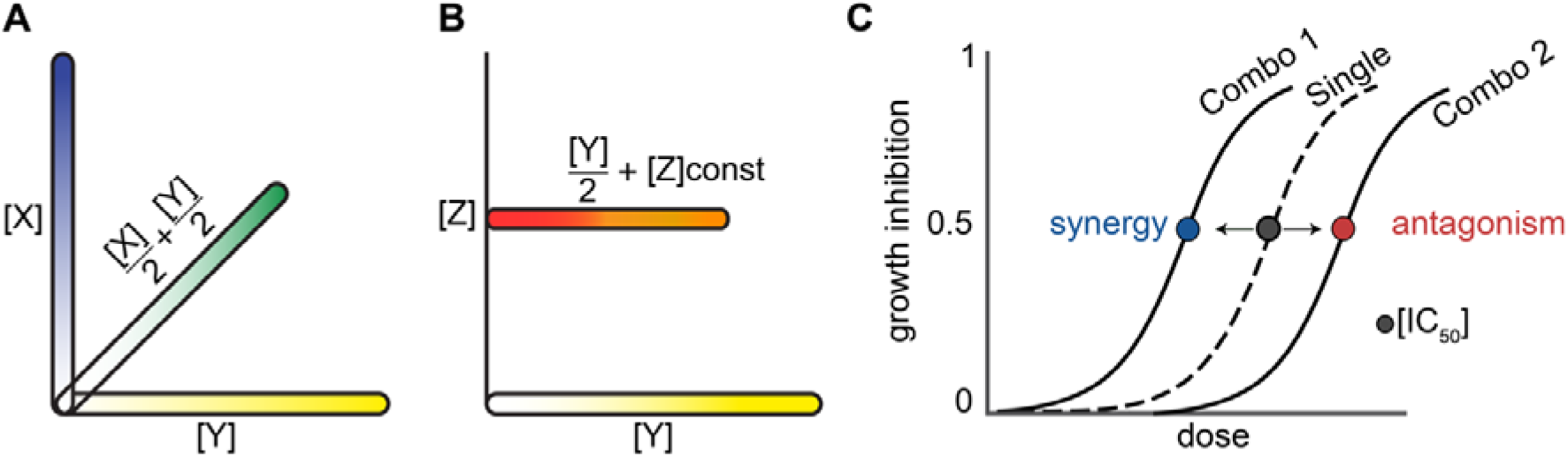
Schematic of drug interaction and potentiation measurement with DiaMOND. (A) DiaMOND single (axes) and pairwise (diagonal) dose-response sampling using an equipotent mixture of two drugs. The x- and y-axes show the doses of the single drugs (X and Y) sampled from low to high concentrations, as indicated by the blue and yellow gradient. The diagonal is the mixture of half of each single drug at each dose, as shown by the green gradient. (B) Schematic of antibiotic and antibiotic-potentiator dose-response measurement. A dose-response for the antibiotic alone (Y; yellow gradient) and with the addition of a constant concentration of the potentiating agent (Z; red to orange gradient). The antibiotic-potentiator dose-response is a mixture of half the dose of the potentiating drug (constant) and half of the non-potentiating drug (increasing amounts in a dose-response). (C) Schematic of shifts in dose-response curves with synergy or antagonism. The dose-response curve in the dashed line represents the effect of a single drug. When the single drug is combined with another drug, the combination curve might shift to the left, indicating synergy (blue dot), or shift to the right indicating antagonism (red dot).

Another complementary approach is to design combinations that include potentiators, e.g., inactive compounds as single agents that increase the efficacy of other drugs. Similarly, an inactive drug can also decrease the efficacy of other drugs, and we refer to this effect as attenuation. This approach could be particularly relevant in designing combination therapy against *M. abscessus*, where there is a dearth of potent antibiotics for clinicians to choose from. For example, SPR741, a synthetic polymyxin analog with little effective against gram-negative bacteria as a stand-alone agent, exhibits synergy in combination with other antibiotics (20–22). In *M. abscessus*, verapamil has minimal activity against ATCC19977 but potentiates the activity of bedaquiline (23). Another reason to search for potentiator-antibiotic pairs is the possibility of reducing treatment doses to alleviate side effects. For instance, intravenous amikacin can cause adverse effects, including gastrointestinal distress (e.g. nausea) and serious cases of ototoxicity and nephrotoxicity (24). A partner drug to amikacin that potentiates its activity could lower the treatment dose and reduce adverse effects.

DiaMOND was initially designed to quantify drug interaction between two potent drugs. Therefore, modification is required to quantify drug interaction between an active drug and a potentiator candidate. Here, we defined potentiator candidates as compounds that did not achieve growth inhibition at a clinically achievable concentration in patients without extreme adverse side effects. We have included four potentiator candidates in our study: avibactam, tetracycline, streptomycin, and verapamil (Table 1). Because potentiator candidates are not active as single drugs, we developed a geometrically optimized sampling of the checkerboard assay to measure the effect of the potentiator candidate at a constant dose on the potency of effective (dose-responsive) antibiotics (Fig 1B). We measured two-dose responses: (a) the single-drug dose-response curve [Y] with increasing concentrations and (b) the single-drug dose-response [Y] (in which drug concentration was reduced by half for all doses) combined with a fixed-dose (1.5x reported maximum plasma concentration after 12-24 hours of dosing in humans) of drug [Z] (Fig. 1B) (25). The effect of the potentiator candidate was calculated as a fold change in concentration of the antibiotic to reach a specific level of growth inhibition (IC_50_ or IC_90_) with the potentiator candidate compared to the drug alone. This fold shift in IC (FsIC) ratio can be interpreted similarly to FIC values: log_2_FsICs are negative for potentiator-drug pairs and positive for attenuator-drug pairs. We focused on evaluating fold shifts at 50% and 90% growth inhibition levels (FsIC_50_ and FsIC_90_, respectively). Potentiation and attenuation can be visualized by the shift of combination dose-response curves relative to single-drug dose-response curves resembling synergy and antagonism, respectively (Fig.1C), with the exception that the combination dose-response utilizes a constant dose of the potentiator candidate with an increasing dose level of the antibiotic.

### A drug interaction landscape of ATCC19977

To generate a systematic dataset of drug interactions for *M. abscessus*, we measured 153 pairwise combinations of 18 drugs representing five drug classes in the reference strain ATCC19977 (Fig. 2A). Among these combinations, we report 122 combinations that passed our quality control metrics. The details of quality control metrics can be found in the method section. Briefly, passing criteria include metrics such as Z-factor, the quality of the dose-response curve fitting, and data reproducibility. The most common reason for quality control failure was variation in potency that compromised the equipotent design of the combination dose-response curve, leading to poor equipotency score and reproducibility. Interactions were measured up to nine times; combinations that failed to achieve at least biological duplicates that passed these criteria were categorized as unmeasurable and reported as N/A (gray boxes in Fig. 2A) in this dataset. Approximately 20% of the drug interactions were unmeasurable due to extreme variation in drug combination response, which is consistent with challenges in measuring drug susceptibility to *M. abscessus* due to lack of reproducibility (26). Nonetheless, it is important to understand which drug combinations cannot be reliably measured.

**FIG 2.**
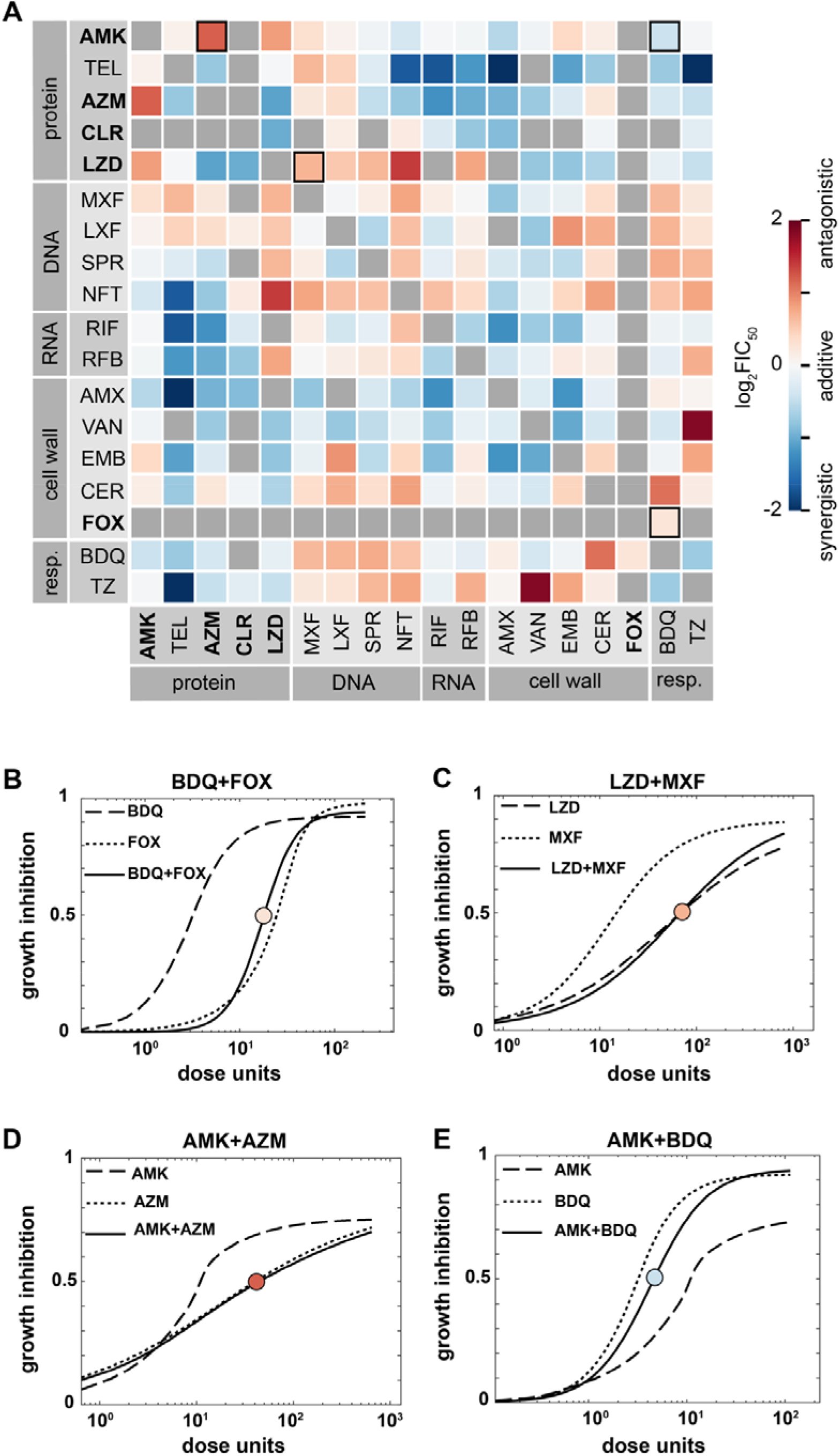
Drug interaction landscape of *M. abscessus strain* ATCC19977. (A) Heatmap of pairwise drug interactions among 18 drugs. Drugs are organized by the mechanism of action, and drugs recommended for treating *M. abscessus* infection are indicated in bold text. Drug interactions are evaluated with log_2_FIC_50_ values: log_2_FIC < 0 (synergy, blue) and log_2_FIC > 0 (antagonism, red). Gray boxes indicate unmeasurable drug combinations due to poor reproducibility. Outlined squares indicate combinations that are shown in the dose responses below. (B-E) Example pairwise (and corresponding single-drug) dose responses. Pairwise dose responses for synergistic and antagonistic combinations are shifted to the left and right, respectively, as compared to the corresponding single-drug dose-response curves. Circles with gray indicate the IC_50_ of single drugs, and circles with the red-blue scale according to log_2_FIC_50_ values as in (A). The y-axis represents growth inhibition, whereas the x-axis represents the dose unit (a unitless representation of the volume, or concentration, used for each drug. A dose unit is preferred when plotting as different drugs have different inhibition concentrations, while a dose unit normalizes the difference and allows easy representation) (B-C) Dose-response curves of combinations that were reported from other studies which are bedaquiline (BDQ) + cefoxitin (FOX) (B) and linezolid (LZD) + moxifloxacin (MXF) (C). (D) Dose-response curves for clinically relevant combination amikacin (AMK) + azithromycin (AZM) (antagonistic) and (E) novel combination amikacin (AMK) + bedaquiline (BDQ) (synergistic).

To understand how well DiaMOND drug interaction measurements correlate with other independent studies, we compared our measurements with previously reported results. We observed a qualitative (synergy vs. antagonism) agreement between drug interactions from DiaMOND and other studies. For example, bedaquiline has been shown to eliminate the bactericidal effect of ß-lactam by dampening the overproduction of ATP to toxic levels normally found upon treatment with these drugs (27). Using DiaMOND, we observed a mild antagonism (log_2_FIC_50_ of 0.25) between bedaquiline and cefoxitin, a ß-lactam (Fig. 2B). The antagonism can be visually observed by the right shift of the combination dose-response curve (solid line), compare to the expected combination dose-response curve if ½ concentration of bedaquiline and ½ concentration of cefoxitin were used (not shown in the figure). Similarly, we observed a mild antagonism between linezolid and moxifloxacin in agreement interactions reported in Zhang et al. (log_2_FIC_50_ of 0.67 and 0.17, respectively, Fig. 2C) (28). Combinations such as rifabutin and clarithromycin were reported to be synergistic by Pyrjma et al. (29). Using DiaMOND, we also observed that rifabutin and clarithromycin are synergistic (log_2_FIC_50_ of −0.76 and −3.1, respectively). Taken together, we conclude that DiaMOND measurements of *M. abscessus* drug interactions are comparable to traditional checkerboard approaches previously reported in other studies.

In other bacterial species, including *M. tuberculosis* and *E. coli*, drug interactions tend toward antagonism (30). In our study, we observed a tendency toward synergy in the landscape of drug interactions in *M. abscessus*. Of the 122 measurable pairwise drug interactions among 18 drugs, about two-thirds (71) were synergistic or additive (with ~ 0 and negative log_2_FIC_50_ values). Our data suggests that there is potential to improve *M. abscessus* multi-drug regimens using combinations of existing antibiotics. To further explore these combinations, we analyzed our dataset in smaller sections for interpretability.

#### Amikacin and macrolides (azithromycin and clarithromycin)

Combination treatment with amikacin and azithromycin (or clarithromycin) is a recommended treatment for macrolide-susceptible *M. abscessus* (12). We observed a strong antagonistic relationship between amikacin and a commonly used macrolide, azithromycin (Fig. 2D, log_2_FIC_50_ of 1.2). We also tested another macrolide-amikacin pair (clarithromycin-amikacin) but found that the combination was unmeasurable because the data fluctuated significantly from replicate-to-replicate. Generally, we did not observe any strong synergistic combinations between amikacin and other tested drugs, except with amoxicillin, bedaquiline, and nitrofurantoin (log_2_FIC_50_ of −0.58, −0.43, and −0.36, respectively). None of these drugs are currently being recommended to be used with amikacin. For azithromycin and clarithromycin, we found more synergistic combinations between these macrolides and other tested drugs. Both azithromycin and clarithromycin are strongly synergistic with linezolid (log_2_FIC_50_ of −1.1 and −0.99, respectively), an oxazolidinone that is suggested to treat *M. abscessus*. Azithromycin and clarithromycin are also synergistic with cell-wall-acting drugs (except with cerulenin), RNA polymerase inhibitors, and respiratory inhibitors (Fig. 2A). These data suggest that there is potential to improve treatment outcomes by combining these core bacteriostatic protein elongation inhibitors with antibiotics that disrupt initiation of protein synthesis (linezolid) or target other cellular processes.

#### Clinically used antibiotics and other common antituberculosis agents

Besides amikacin and macrolides, other drugs recommended for treating *M. abscessus* are imipenem, cefoxitin, tigecycline, clofazimine, and linezolid (12). Among this list, we focused on linezolid and cefoxitin due to drug availability and reproducibility. **Linezolid:** We observed synergies between linezolid and some of the drugs used in the clinic, such as azithromycin and clarithromycin (log_2_FIC_50_ of −1.1 and −1.0, respectively). Linezolid is strongly antagonistic with DNA-targeting antibiotics in our drug set while synergistic with cell-wall-acting drugs and respiratory inhibitors (Fig 2A), suggesting an underlying relationship between drug interaction and mechanism of action. We did not observe a synergistic relationship between linezolid and amikacin, which was previously reported in another study (28). However, there are several differences between the two studies that may account for these differences, including the media used (7H9 medium with supplements vs. CAMHB), strains (ATCC19977 vs. clinical isolates from patients), and method used to determine drug interaction (DiaMOND vs. broth microdilution). **Cefoxitin:** Cefoxitin is one of the few ß-lactams recommended to treat *M. abscessus*, besides imipenem (31). We could not obtain a reliable dose-response curve for both imipenem and cefoxitin, which may be due to the instability of ß-lactams in solution or the presence of the blamab gene that confers their resistance to ß-lactam. Nevertheless, one combination that passed our quality control is bedaquiline and cefoxitin (log_2_FIC_50_ of 0.25). The antagonistic relationship between bedaquiline and cefoxitin has been reported in another study (32). It is thought that cefoxitin (and imipenem) triggers an ATP burst in *M. abscessus* by increasing oxidative phosphorylation, which is suppressed by bedaquiline, an F-ATP synthase inhibitor, leading to an elimination of the bactericidal effect of cefoxitin against *M. abscessus* (32, 33). Though this result is preliminary, it suggests that combining ß-lactams with bedaquiline could be unfavorable. **Moxifloxacin:** Moxifloxacin and other fluoroquinolones have been shown to have good activities against *M. abscessus* isolates *in vivo* (34). We and others have observed that moxifloxacin is antagonistic with azithromycin *in vitro* (log_2_FIC_50_ of 0.23) and *in vivo* (35). We also observed that moxifloxacin is antagonistic with recommended antibiotics such as amikacin and linezolid (log_2_FIC_50_ of 0.31 and 0.68 respectively), suggesting its inclusion in multidrug therapies should be carefully considered. **Ethambutol:** Ethambutol is another antituberculosis agent with potential activity against NTMs such as *Mycobacterium avium* complex (MAC) due to its ability to slow the acquisition of macrolide-resistance clinically (36). The interactions of ethambutol with other drugs used to treat NTMs are unknown. Here, we observed a synergistic relationship of ethambutol with linezolid or azithromycin but not amikacin (log_2_FIC_50_ of −0.79, −0.28 and 0.4 respectively), suggesting potential for exploring of ethambutol for *M. abscessus* treatment. **Rifampicin:** Rifamycin is a common antituberculosis agent that is highly potent *in vivo* and reduces relapse rate in combination therapies (37). However, rifampicin is not commonly used for *M. abscessus* because they are known to be intrinsically resistant to rifampicin (38). *M. abscessus* resistance to rifampicin partially due to its intrinsically low cell-wall permeability, possibly due to high lipid content, as well as drug efflux pump (37, 39). Additionally, *M. abscessus* encodes rifampicin ADP-ribosyltransferase (Arr_Mab_), whose function is to catalyze ADP-ribosylation and render rifampicin inactive (40, 41). Though rifampicin does not have high potency against *M. abscessus*, our dataset demostrate synergistic interaction between rifampicin with many tested drugs, including amikacin and azithromycin. Rifampicin also synergizes all cell-wall-acting antibiotics in our dataset; the mechanism of synergy may be due to a weakened cell wall allowing better entry for rifampicin. Though rifabutin, another rifamycin antibiotic, has been approved to treat Mycobacterium Avium Complex and TB, its potential against *M. abscessus* has not been fully explored. Recent findings demonstrate limited modification of rifabutin by the Arr_Mab_ in comparison to rifampicin, resulting in lower MICs against both the ATCC19977 strain and clinical *M. abscessus* isolates (42). We also observed in our dataset that rifabutin has a lower IC_50_ and is more potent than rifampicin (IC_50_ of 1.8μg/mL versus 16 μg/mL, respectively, Table 1, Fig. 4A). However, rifabutin in our pairwise combinations tend toward antagonism, indicated by more red squares in the heat map compared to rifampicin (Fig. 2A). Nevertheless, rifabutin remains synergistic with frontline *M. abscessus* antibiotics such as clarithromycin (which has also been reported by other study), amikacin, and azithromycin, suggesting that rifabutin may be part of a clinically useful combination therapy (43). The rifamycin pairwise interaction data support continued exploration of this important class of drugs for treatment of *M. abscessus* infections.

**FIG 3.**
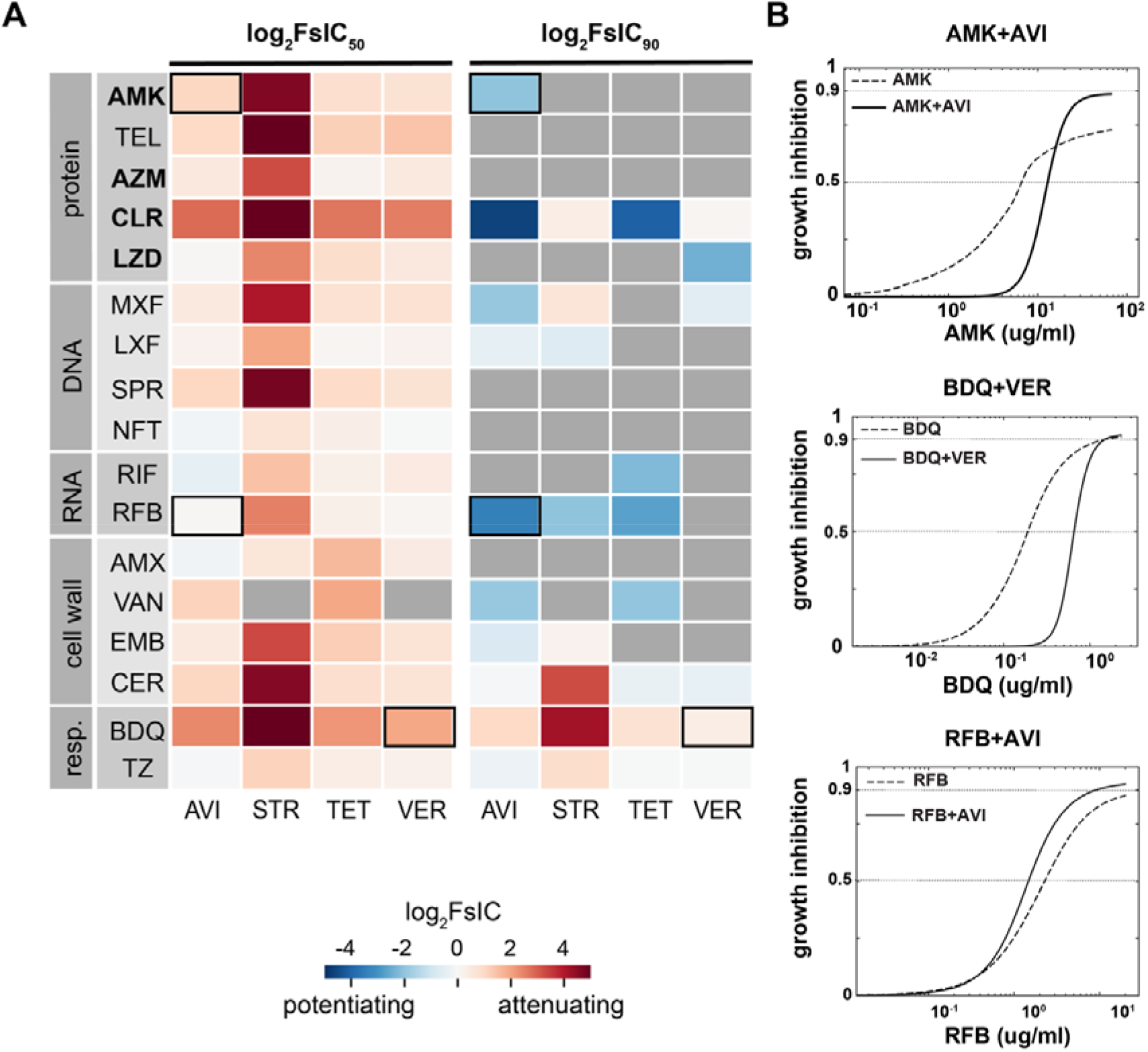
Effect of potentiator candidates on antibiotic efficacies in *M. abscessus* ATCC19977. (A) Heat map of pairwise drug interactions with potentiators at a fixed concentration. Drugs are categorized based on their mechanism of action and drugs recommended for treating *M. abscessus* infection are indicated in bold text. Drug interaction measurement is expressed as the change in log_2_ fold shift at IC_50_ (left) or IC_90_ (right). Log2FsIC < 0 indicates potentiating effects, shown in blue, and log_2_FsIC > 0 indicate attenuating effects, shown in red. Outline squares indicate combinations that are shown in the dose-response below. (B) Example dose-response curves showing the fold shift in IC_50_ and IC_90_ for drugs combined with potentiators, amikacin (AMK) + avibactam (AVI) (top), bedaquiline (BDQ) + verapamil (VER) (middle), rifabutin (RFB) + avibactam (AVI) (bottom). The dash line indicates the single drug dose-response curve, while the solid line represents the combination curve. A left shift in the solid line dose-response curve indicates a potentiating effect, and a right shift shows an attenuated effect by the potentiator.

**FIG 4.**
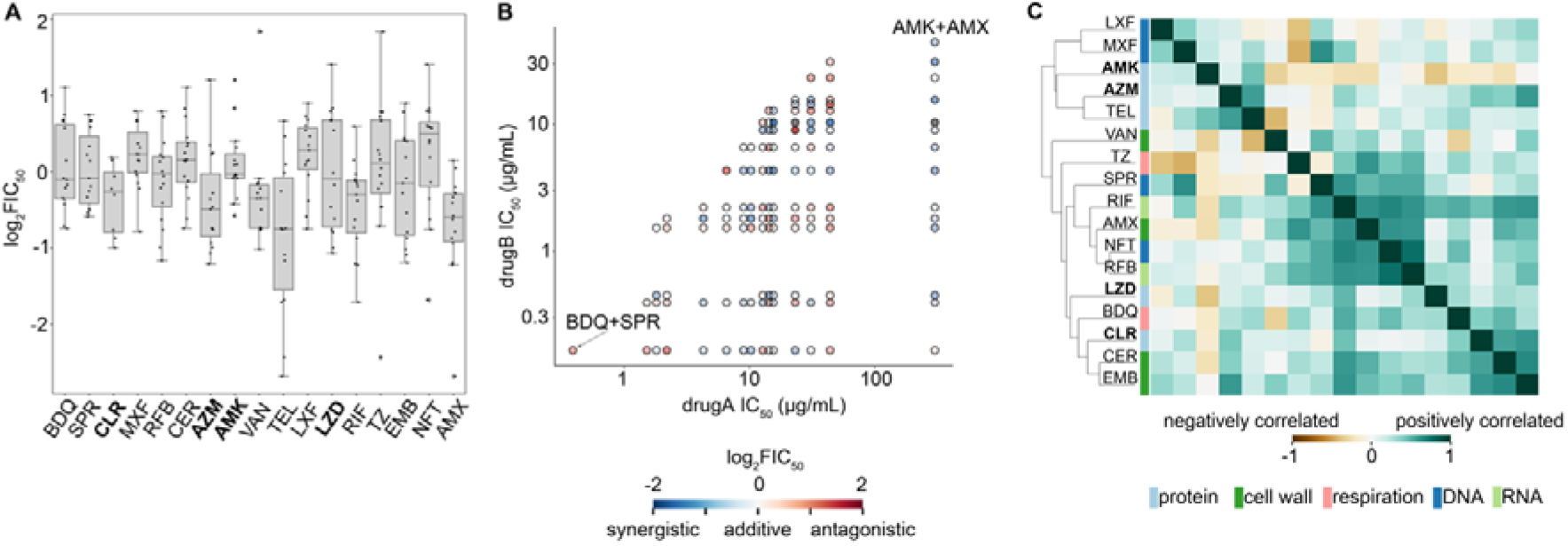
Relationship between drug potency or mechanism of action with synergy. (A) Box plot of drug interaction (log_2_FIC_50_) of all combinations containing the single drug represented on the x-axis, drugs recommended for treating *M. abscessus* infection are indicated in bold text. Each dot inside the box represents a combination of that single drug and another drug in the data set. Single drugs are ordered on the x-axis based on IC_50_ (Table 1), with drugs having the highest IC_50_ on the right and the lowest IC_50_ on the left (B) Comparison of drug interaction scores with single drug potency in ATCC1977. Pairwise combinations are plotted as the components of their single drug IC_50_ (μg/mL). Given a pairwise combination, the potency of a single drug is plotted on the y-axis while the non-potent drug is on the x-axis; drugs recommended for treating *M. abscessus* infection are indicated in bold text. The color fill indicates the synergistic (blue) or antagonistic (red) interaction in ATCC19977. (C) Clustering of drugs’ interaction profile in ATCC19977. Each box represents the Pearson correlation of drug interaction profiles between two antibiotics. 1 indicates a positive correlation, whereas −1 indicates a negative correlation. The color bar along the y-axis represents the general drug target pathway (e.g., inhibition of protein, cell wall, DNA, or RNA synthesis and inhibition of respiration.)

### Non-traditional NTM antibiotics

Next, we evaluated an unexplored space in combination therapies by analyzing drug interactions among antibacterials that are not traditionally used to treat NTMs and TB. **Telithromycin:** Telithromycin belongs to a large group of ketolides, a newer generation of macrolide with a slightly different mechanism of action (44, 45). Ketolides appear to partially inhibit protein synthesis rather than complete or near-complete inhibition of protein synthesis, which leads to more cellular degradation and a stronger bactericidal effect (45, 46). Despite being designed to be an alternative for some macrolide-resistant bacteria, telithromycin is not often used in the clinic due to its toxic side effects on the liver (47). We observed particularly strong synergies between telithromycin and other antibacterial compounds such as nitrofurantoin, rifampicin, amoxicillin, thioridazine, and azithromycin (log_2_FIC_50_ of −1.7, −1.7, −2.7, −2.4, and −0.76, respectively), suggesting the potential of using combinations containing telithromycin to treat macrolide-resistant *M. abscessus*. Nevertheless, telithromycin is mildly antagonistic with amikacin and additive with linezolid (log_2_FIC_50_ of 0.09 and −0.02, respectively). Therefore, further evaluation may be required before combining telithromycin with these clinically recommended drugs. **Amoxicillin and vancomycin:** Cell-wall-acting antibiotics amoxicillin and vancomycin show broad synergies with different classes of antimycobacterial (Fig. 2A). We found that amoxicillin is synergistic with almost all the tested partner drugs, including amikacin, azithromycin, clarithromycin, rifampicin, and ethambutol (log_2_FIC_50_ of −0.58, −0.94, −0.87, −1.2, and −1.2, respectively). Amoxicillin has previously been known to enhance the effect of ß-lactam antibiotics such as imipenem and relebactam (48). Our results suggest that amoxicillin may also partner well with a broad range of antibiotics. For example, amoxicillin and telithromycin are strongly synergistic, with a log_2_FIC_50_ of −2.7 (the most synergistic combo in our dataset). Vancomycin is another widely synergistic drug in our drug panel and was reported to be synergistic with clarithromycin, one of the most used clinical drugs for *M. abscessus* (49). Although we did not obtain a reliable drug interaction measurement for vancomycin with clarithromycin due to poor reproducibility, vancomycin is a promising candidate for a multi-drug regimen which showed broad synergy with most of the other drugs tested, except with thioridazine (Fig. 2A). Together, these data suggest that telithromycin, vancomycin, and amoxicillin, despite being uncommon drugs to choose in the clinic, tend toward synergy and may be candidates as partner drugs for *M. abscesses* regimen development.

### Bedaquiline

Another drug candidate for the combination treatment of *M. abscessus* is bedaquiline, a diarylquinoline that inhibits subunit c of mycobacterial ATP synthase. In a zebrafish model of *M. abscessus* infection, bedaquiline was shown to exert a therapeutic effect by preventing the formation of abscesses (50). Additionally, a preliminary study in the clinic demonstrated the efficacy of using bedaquiline as salvage therapy. Within three months of treatment, bedaquiline reduced the bacterial load in the sputum of patients (51). Consistent with this finding, we found bedaquiline to have a low IC_50_ value of 0.16 μg/mL (Table 1). With DiaMOND, we observe that bedaquiline mildly antagonizes cefoxitin and amoxicillin (log_2_FIC_50_ of 0.25 and 0.14, respectively) and is mildly synergistic with vancomycin and ethambutol (log_2_FIC_50_ of −0.35 and −0.08. respectively). The antagonistic relationship between bedaquiline and cefoxitin in our data set is in agreement with previous studies that suggested bedaquiline eliminates the effects of ß-lactams (52). To evaluate the potential of bedaquiline more fully, we investigated drug interactions between bedaquiline and other antibiotics where these drug interactions have not yet been reported. We observed an overall trend that bedaquiline is antagonistic with DNA-acting antibiotics used in this study (Fig. 2A). In contrast, bedaquiline is synergistic with protein synthesis targeting drugs and thioridazine (Fig. 2A). Bedaquiline’s potency and synergy with clinically favored protein synthesis inhibitors, such as amikacin and azithromycin (log_2_FIC_50_ of −0.42 and −0.35, respectively) have not been previously reported to our knowledge. However, the administration of bedaquiline following amikacin yields positive clinical outcomes and has been reported for macrolide-resistant *M. fortuitum* complex soft tissue infection (53). A small preliminary report also suggested the potential clinical and microbiologic activity of bedaquiline against *M. abscessus* (51). Though current clinical results on bedaquiline are limited and further studies are required, these results suggested that bedaquiline should be explored for treating *M. abscessus*, and combinations containing bedaquiline could be considered for macrolide-resistant cases.

**Thioridazine** is an atypical antipsychotic still available in generic form, but with limited clinical use because of QTc prolongation (54). This agent initially gained interest for its potential use in treatment of MDR-Tb, as it has been shown to have properties against Mtb *in vitro*, *in vivo*, and in clinical cases, irrespective of antibiotic resistance status (55). In this study, thioridazine is synergistic with clinically preferred drugs such as amikacin, azithromycin, and clarithromycin (log_2_FIC_50_ of −0.04, −0.46, and −0.22, respectively), as well as drugs that are not traditionally used for NTM treatment such as telithromycin, and bedaquiline (log_2_FIC_50_ of −2.4 and −0.72, respectively). Although thioridazine does not appear to have a strongly synergistic drug interaction profile (Fig. 2A), it is one of a few drugs in this dataset that shows synergy with clinically preferred drugs (other drugs are bedaquiline, vancomycin, and amoxicillin), suggesting that non-traditional antibiotics such as thioridazine could be useful in combination with other agents for *M. abscessus* infection, where the intrinsic resistance to common antibiotics is a challenge for treatment success.

### SPR719

Another promising candidate for *M. abscessus* multi-drug regimen development is SPR719. SPR719 is a DNA gyrase inhibitor (GyrB) that is currently in clinical studies as a treatment for NTM infections, and is the prodrug version of SPR720, which is in phase I clinical development as a new oral agent for NTM infection and Mtb infection (56). We determined the IC_50_ of SPR719 to be 0.39 μg/mL (Table 1), in agreement with the reported IC_50_ from previous studies (0.25 – 4 μg/mL, (26)). Despite being highly potent, the interaction profile of SPR719 is relatively mild, with a balanced profile of synergies and antagonisms (an even number of synergistic combinations, having log_2_FIC_50_ < 0 and antagonistic combinations log_2_FIC_50_ > 0 (Fig. 2A)). SPR719 is additive with amikacin (log_2_FIC_50_ of −0.09) and synergistic with drugs from a broad range of classes used to treat NTMs and TB, including azithromycin, telithromycin, rifampicin (log_2_FIC_50_ of −0.49, −0.26 and −0.19, respectively), and all cell-wall acting antibiotics tested (Fig. 2A). Through genomic analysis, *M. abscessus* was shown to contain a natural A92S mutation in *gyrB*, which is unique among other NTMs and *M. tuberculosis* (57). This mutation causes a conformational change in the ATP binding site, which may partially explain the intrinsic resistance of NTMs to GyrB inhibitors (57, 58). Despite this mutation, SPR719 is still very potent against *M. abscessus* (Table 1, Fig. 4A) (58). SPR719 is more potent than the fluoroquinolone moxifloxacin and is synergistic with more combinations compared to moxifloxacin (moxifloxacin log_2_FIC_50_ median is 0.22 with only four combinations having log_2_FIC_50_ < 0) (Table 1, Fig. 4A). Although moxifloxacin is not commonly recommended to treat *M. abscessus*, moxifloxacin is still being used in conjunction with amikacin and macrolides in clinical settings (12, 59). Together, these data suggest that SPR719 may be part of improved combination therapies for *M. abscessus*, including as a replacement for moxifloxacin.

### Antibiotic potentiation to improve combinations therapy against *M. abscessus*

To determine whether a potentiator candidate enhances or attenuates the potency of antibiotics, we measured how the addition of potentiator candidates shifts the partner antibiotic’s dose-response curve (Fig. 1B-C, Fig. 3A). We quantified the degree of potentiation (and attenuation) by calculating the log_2_-transformed fold shift in IC_50_ or IC_90_ of the effective antibiotic with the addition of a constant dose of the candidate potentiator so that a negative log_2_FsIC indicates potentiation and a positive value indications attenuation (Fig. 1B-C). We observed that all tested antibiotics were attenuated by all four potentiator candidates at IC_50_, but some antibiotics were potentiated at IC_90_ (Fig. 3A). For example, avibactam attenuates the effect of amikacin at IC_50_, indicated by the right shift in combination dose-response curves compared to the amikacin dose-response curve (log_2_FsIC_50_ of 1.1, Fig. 3B). However, with avibactam, amikacin reached IC_90_ at a lower concentration, indicated by a left shift in the combination dose-response curve compared to single-drug dose-response curve (log_2_FsIC_90_ of −2.1, Fig. 3B), indicating a potentiating effect. We observe a similar shift in all combinations where IC_90_ is reported, e.g., potentiator candidates shifted from being attenuating to either less attenuating (such as bedaquiline with streptomycin; log_2_FsIC shifts from 6.4 at IC_50_ to 4.2 at IC_90_), or from attenuating to potentiating (as with the previous example between amikacin and avibactam), or from less potentiating to more potentiating (such as avibactam and rifabutin; log_2_FsIC shift from 0.04 at IC_50_ to −3.4 at IC_90_, Fig. 3B). This dramatic shift from attenuation to potentiation is explained by an increase in the steepness of the dose-response curve with the addition of potentiator candidates so that the effect in some combinations is an increase in potency at higher dose levels of the antibiotic (Fig. 3B). Among four tested potentiator candidates, streptomycin has the strongest attenuating effect at IC_50_ and IC_90_ (except with rifabutin and levofloxacin, in which case streptomycin acts as a potentiator). Among our tested drug set, three inactive drugs potentiate the activity of rifabutin (with the 4^th^ being unmeasurable).

Beta-lactams are the most widely used antibiotics (60). However, due to the presence of the broad-spectrum beta-lactamase BlaMab, only imipenem and cefoxitin are currently recommended in the multidrug-regimens targeting *M. abscessus* (8). Therefore, potentiators such as avibactam, a non-ß-lactam ß-lactamase inhibitor shown to efficiently inhibit BlaMab, can potentially help extend the spectrum of ß-lactam antibiotics active against *M. abscessus* (61). Porins of *M. abscessus* cell wall have been shown to partially contribute to ß-lactam resistance as they allow hydrophilic molecules to cross the cell membrane, which can interact with the target in the cytoplasm and potentially trigger the expression of resistance genes (10, 62). Avibactam has also been reported to reduce the MIC of cell-wall-acting agents (63), and we observed a potentiating effect of avibactam with cell-wall-acting agents ethambutol, cerulenin, and vancomycin (log_2_FsIC_90_ of −0.76, −0.072, and −1.9) at IC_90_. Because avibactam is a ß-lactamase inhibitor, we also wanted to determine its effect with amoxicillin. Due to the low potency of amoxicillin, there is no IC_90_ for amoxicillin. However, at IC_50_, we observed a mild potentiating effect (log_2_FsIC_50_ of −0.2), which is not commonly observed in our dataset (the other combinations are avibactam and nitrofurantoin, log_2_FsIC_50_ of −0.2, avibactam and rifampicin, log_2_FsIC_50_ of −0.44, and avibactam with thioridazine, log_2_FsIC_50_ of −0.061).

Although verapamil is not effective on its own and have limited clinical used due to its primary effect on cardiac function, verapamil at 50 μg/mL was reported to enhance the killing of bedaquiline in *M. abscessus* (23, 64). In contrast, our data show a right shift (attenuation) of the bedaquiline + verapamil dose-response curve compared to the bedaquiline single dose-response curve (log_2_FsIC_50_ value of 1.9, Fig. 3B). This difference in the role of verapamil on bedaquiline efficacy may be due to experimental differences such as verapamil concentration. In Viljoen et al., 50μg/mL of verapamil was used, whereas we used a lower dose of 2.35 μg/mL, which was chosen based on 3-fold maximum plasma concentration 4-6 hours after administration, assuming one pill of 100 mg of bedaquiline was administered (65).

We observed that streptomycin and verapamil largely attenuated the potency of partner drugs, whereas avibactam and tetracycline are broadly potentiating at IC_90_ (Fig. 3A), suggesting that these two candidates should be further explored as an element of multi-drug treatment for *M. abscessus* infection. Additionally, we showed that with modification, DiaMOND can be used to measure the combined effect between a potentiator and an active drug.

### Drug interactions cannot be predicted from single drug potencies or mechanisms of action

In the absence of a known effective multidrug therapy for *M. abscessus* infection, clinicians usually rely on single drug susceptibility profiles and their own experience to determine the best drug combination for each patient. In some cancers, single-drug susceptibility profiles can be used to design optimized combination therapies under the principle that efficacy for each cancer is determined by its susceptibility to any of the agents and that combination therapies were effective as bet-hedging strategies (66). To understand whether we could predict drug interactions based on single-drug properties, we started by evaluating whether single-drug potencies were correlated with the propensity for drug interactions to be synergistic or antagonistic in *M. abscessus*. In Fig 4A, single drugs are organized based on their IC_50_ on the x-axis, and the y-axis shows the distribution of log_2_FIC_50_ of all combinations containing that single drug. We do not observe a correlation between distributions of log_2_FIC_50_ compared to IC_50_ for each drug, suggesting that drug interactions cannot be predicted from single drug potencies. We wondered if the most potent or least potent antibiotic is the driver of drug interaction, which may be obscured by looking at the overall propensity for synergy compared to single-drug potencies. To take the IC_50_ of both drugs in each pairwise combination into consideration, we evaluated whether there were patterns of synergy and antagonism compared to the IC_50_s of both partner drugs (Fig. 4B). In the drug interactions among 18 antibiotics, there was no clear trend of synergy and antagonism based on single-drug potencies (e.g., the drug interactions, colored by log_2_FIC_50_, are not clustered by drug potency to either drug). For example, SPR719 and bedaquiline are highly potent drugs with low IC_50_ (IC_50_ of 0.39 μg/mL and 0.16 μg/mL, respectively), but the combination between SPR719 and bedaquiline are antagonistic (log_2_FIC_50_ of 0.74). Conversely, amikacin and amoxicillin are not as potent as SPR719 or bedaquiline (IC_50_ of 6.6 μg/mL and 300 μg/mL, respectively) but are mildly synergistic in combination (log_2_FIC_50_ of −0.59). Together, our analysis demonstrates that single drug potency is not predictive of drug interactions, and we cannot anticipate whether a drug pair will be synergistic or antagonistic based on their potency profiles as single agents.

Next, we evaluated whether there were drug interaction patterns that correspond to each partner drug’s mechanism of action. To test the hypothesis that drugs targeting similar pathways would have similar interaction profiles, we compared the similarities among drug interaction profiles from the 17 antibiotics that target four general processes using hierarchical clustering (Fig. 4C). Cefoxitin is removed from this analysis because there is only one reportable combination containing cefoxitin in our dataset. In general, it is unclear whether drug interaction profiles are clustered by the mechanism of action. For example, rifabutin and nitrofurantoin have similar interaction profiles but different mechanisms of action. However, there are also examples of drugs with a similar mechanism of action that have similar interaction profiles, such as cell-wall acting agents ethambutol and cerulenin, the fluoroquinolones levofloxacin and moxifloxacin, protein synthesis inhibitors azithromycin and telithromycin (Fig. 4C). Nevertheless, it is possible that an analysis of drug interaction profiles across a larger set of drugs may reveal patterns that are not present in this dataset due to the small number of representatives for each drug class. Further analysis with a larger drug set with an even greater number of representatives for each drug category may be necessary to understand whether drug interaction patterns are similar in *M. abscessus* for drugs with closely related mechanisms of action. However, our data do not suggest that there are strong similarities in drug interactions for drugs that target the same pathways.

### Drug interaction is strain-specific

An extensive study of 85 clinical isolates from *M. abscessus* subspecies demonstrated species-specific drug susceptibility, including clinically favored drugs, such as amikacin and macrolide clarithromycin (67). For this reason, we speculated that drug interactions would also vary from isolate to isolate. To test this hypothesis, we selected three strains that were chosen to represent major types of variation among strains, including colony morphology (e.g., smooth versus rough phenotype) and growth rate (e.g., slow-versus fast-growing). The fastest and the slowest growing strain were the lab reference strain ATCC19977 and TMC2 (4hr and 20hr doubling, respectively, Table S2). ATCC19977 and TMC3 shared similar smooth morphologies, and TMC1 and TMC2 shared similar rough morphologies. Detailed information about the doubling time and phenotype of these clinical isolates is included in Table S2.

We used a focused drug set for isolate-to-isolate comparison; the set includes antibiotics that are recommended for current therapies for NTMs and TB (amikacin, azithromycin, cefoxitin, clarithromycin, moxifloxacin, and rifampicin), in development (SPR719 and bedaquiline), or are broadly synergistic in ATCC19977 (telithromycin). Consistent with previous studies, we observed variation in IC_50_ values from isolate-to-isolate (Fig. 5A) (11, 67). Though susceptibility among strains is also variable for newer drugs, some relative potencies for single-antibiotics are retained across strains. For example, bedaquiline and SPR719 have the highest potency (e.g., lowest IC_50_) for the lab reference strain, and this observation holds for other strains (except for SPR719 in TMC3 strains, Fig. 5A). To understand if colony morphotype or growth rate was correlated with drug susceptibility, we calculated Pearson correlations of IC_50_ values between each strain (Fig. S4A). In general, we observed poor correlations in single-drug susceptibility patterns between strains, even in strains with common morphologies and similar growth rates (ATCC1977, TMC3) and (Table 2, Fig. S4A). The only significant correlation (R = 0.88) observed is between TMC2 and ATCC19977, which differ in both morphology and growth rate (Table 2). Together, our data suggest that isolate-to-isolate differences in drug response are not well correlated with colony morphology and doubling times.

**Table 2.** Characterization of lab strain (ATCC19977) and clinical isolates TMC1, TMC2 and TMC3.

| strain | doubling time (hour) | morphology |
| --- | --- | --- |
| ATCC19977 | 4 | smooth |
| TMC 1 | 12 | rough |
| TMC 2 | 20 | rough |
| TMC 3 | 8 | smooth |

**FIG 5.**
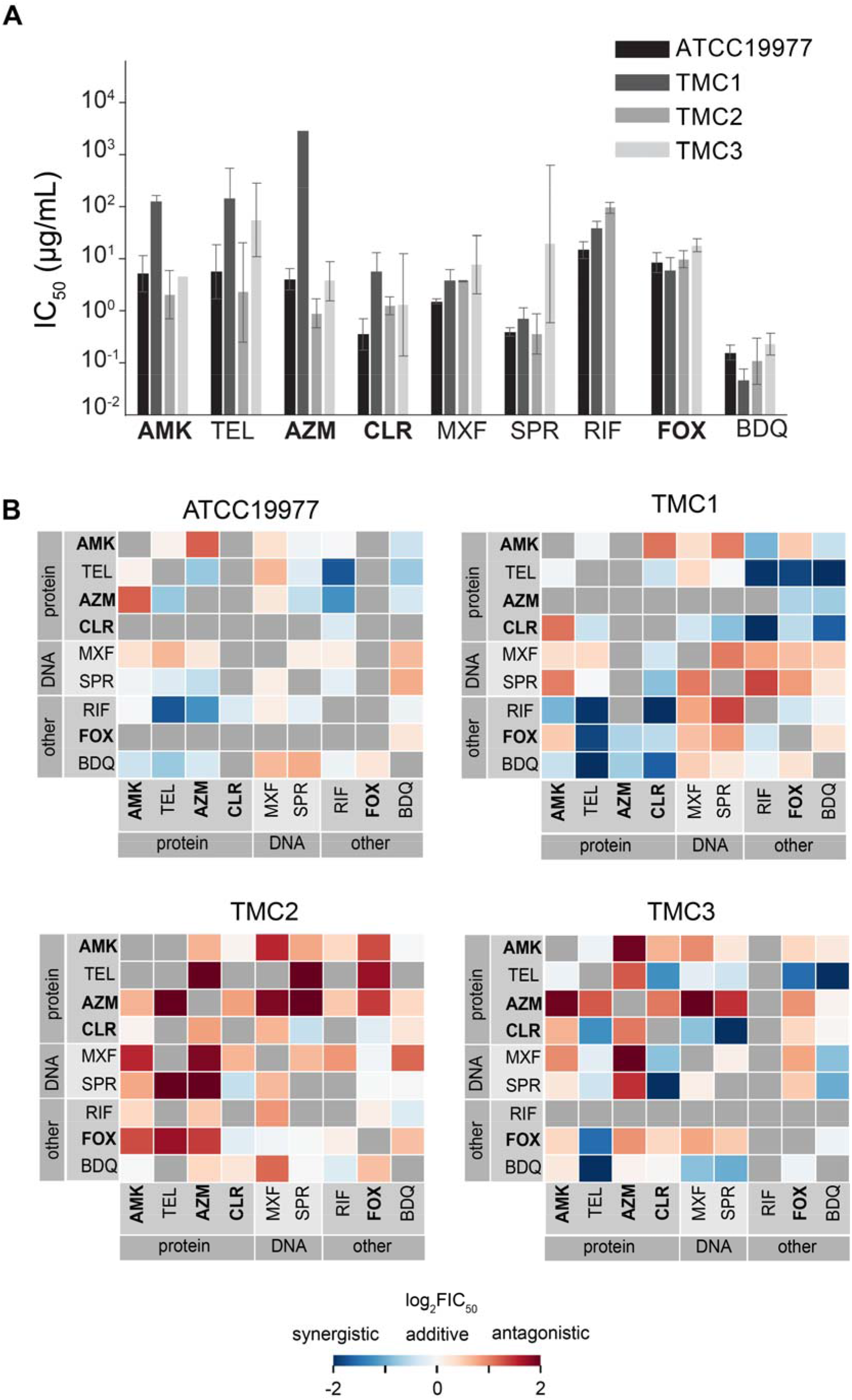
Single drug susceptibility and drug interaction outcomes and in different *M. abscessus* clinical isolates. (A) Single drug susceptibilities for the reference strain (ATCC19977) and three clinical isolates (TMC1, TMC2, and TMC3). Susceptibility is measured by the IC_50_ (μg/ml). Error bars represent standard deviation as compared to the mean, when there are more than one replicates IC values are reported. (B) Heat map showing drug interactions for each strain: ATCC19977 (top left), TMC1 (top right), TMC2 (bottom left) and TMC3 (bottom right). As in Fig 2, drugs are organized by the mechanism of action, and drugs recommended for treating *M. abscessus* infection are indicated in bold text. Drug measurement is expressed in log_2_FIC_50_, where blue highlights log_2_FIC < 0 (synergy) and red is log_2_FIC > 0 (antagonism).

To understand whether drug interactions were also strain-dependent, we measured pairwise drug interactions among the nine drugs in our focused antibiotics set with DiaMOND. The drug interaction profiles varied from isolate to isolate (Fig. 5B). Correlation analysis demonstrates that the most similarity in drug interactions is between TMC1 and ATCC19977 (R = 0.48, Fig. S4B). We noted that there was no correlation in single-drug IC_50_ values between TMC1 and ATCC19977, nor did these two strains share morphologies or growth rates. The lack of similarity between these strains, except in the drug interaction profile, supports our hypothesis that drug interaction is independent of single drug potency and morphology. The spectrum of drug interactions is balanced in ATCC19977, TMC1, and TMC3, whereas TMC2 strongly tends toward antagonism (Fig. 5B). Despite poor overall correlations in drug interaction profiles from strain to strain, we observed some similarities between these strains. For example, amikacin paired with macrolides (such as azithromycin or clarithromycin) is consistently antagonistic. In contrast, combinations of clarithromycin and SPR719 or bedaquiline and telithromycin were all synergistic (Fig. 5B). There are strongly synergistic combinations such as clarithromycin with almost all tested drugs for TMC1 (except amikacin) or telithromycin with almost all tested drugs for TMC3 (except amikacin and azithromycin) (Fig. 5B), suggesting that there may be space for improvement in combination therapy in a strain specific manner. Taken together, our results affirm the need for targeted drug combination testing and suggest that measurements should be made for each isolate for reliable drug interaction determination.

## DISCUSSION

Infections caused by *M. abscessus* are notoriously difficult to treat for many reasons, including innate and acquired drug resistance, low antibiotic efficacies, and variation among strains. Combination antibiotic treatment has the potential to improve treatment efficacies via synergies and potentiation. Despite this potential and reports of several synergistic drug combinations, we have lacked a comprehensive drug interaction dataset to develop drug combinations for the treatment of *M. abscessus*. Here, we applied DiaMOND to systematically measure pairwise drug interactions among 18 drugs against *M. abscessus* to better understand the landscape and potential of combination therapy for effective treatment. Our results include 122 drug pairs for which we report interactions for the first time. These data revealed an unexpectedly large synergy space compared to other bacterial species. We observe many synergistic drug pairs that are currently not considered as options for treating *M. abscessus* pulmonary infection in clinical practice. One example is amoxicillin, which is not currently recommended to be used to treat *M. abscessus*. Amoxicillin is synergistic with almost all tested drugs, combined with its clinical practicality suggests its potential to be used in combination therapy. We did not find that clinically used drug pairs were necessarily synergistic *in vitro*. For example, amikacin and azithromycin is antagonistic *in vitro* while another pair of protein synthesis inhibitors (azithromycin and linezolid) is synergistic. Together, the large number of synergies identified here *in vitro* motivates future investigation of these combinations as a path forward to improve combination therapy to treat pulmonary *M. abscessus* infection.

Our findings demonstrate that some drugs that are not effective alone potentiate the activities of other antibiotics. These agents may play an important role in improving *M. abscessus* regimens given the poor efficacy of antibiotics as monotherapies for *M. abscessus*. To understand whether we could sensitize *M. abscessus* to antibiotics using potentiators, we expanded our combinations screen and adapted the DiaMOND methodology to evaluate how drugs that are considered inactive affect the efficacies of active drugs. Among the four candidate potentiators we tested, avibactam has been subjected to multiple studies due to its ability to inhibit Bla_mab_ and potentially allow more ß-lactams to be used to treat *M. abscessus*. Our results are consistent with previously reported studies on combining a non-ß-lactam ß-lactamase inhibitor with ß-lactam to extend the efficacy of ß-lactam, which suggests that avibactam should be considered in a multi-drug therapy (61, 63). Additionally, the most potentiating effect is observed at 90% growth inhibition (not 50% growth inhibition), suggesting that the potentiators’ effect is likely to be more prominent at higher concentrations of the active antibiotic.

We found that we could not predict drug interactions based on drug potencies or mechanisms of action. Though drug interaction profiles were not very similar among antimicrobials targeting the same cellular process (cell wall, DNA, etc.), we observed that some antibiotics were consistently synergistic or antagonistic with drugs from the same class. For example, DNA-targeting drugs are antagonistic with linezolid and respiratory inhibitors (bedaquiline and thioridazine) and synergistic with cell-wall-acting antibiotics (ethambutol, vancomycin, and amoxicillin). The reason for the observed synergy between cell-wall-acting and DNA-acting compounds may be due to cell wall targeting compounds increasing the permeability of the mycobacterial cell wall, resulting in an increased accumulation of DNA-targeting compounds in the cell (68). *M. abscessus* is known to increase the activity of efflux pump inhibitors when exposed to bedaquiline, which may result in the rapid efflux of DNA-acting compounds and an antagonist interaction (23).

Another challenge with developing an effective regimen for *M. abscessus* is the remarkable variation in drug response among strains, likely due to genomic diversity in NTM species, even across morphologically related isolates (26, 69). We measured drug interactions in three clinical isolates with different colony morphologies and growth rates. Previous studies have shown that the impact of colony morphotype, which includes the transition of the smooth colony to the rough colony during infection, highly affects drug susceptibility (70). To understand whether a universal multidrug regimen may be developed that is effective against all *M. abscessus* strains, we measured a core set of drug pairs in different clinical strains of *M. abscessus*. We found that drug interactions were poorly correlated across strains and could not be systematically determined based on data from the reference strain, strains with similar morphologies or growth rates, the single-drug susceptibility profiles, or the drug mechanisms of action. This result suggests the potential need to include susceptibility testing for drug combinations, instead of single drugs only in clinical laboratory testing. Strain-to-strain variation in drug interaction profile may be determined by subspecies or genetic differences, for instance the presence of the *erm* gene, which can confer macrolide resistance. Future work measuring drug interactions across a large set of clinical isolates is required to address this question. Nevertheless, our observations drug interactions in *M. abscessus* are strain-specific supports the idea that treatment regimen development may need to be personalized for each isolate (e.g., directly measured) rather than derived from guidelines based on other strains.

This systematic study of the drug combination landscape in *M. abscessus* suggests that we require a more extensive phenotypic evaluation of drug interactions to develop improved therapies. Additionally, the relationship between *in vitro* single drug susceptibility or combination measurement and clinical outcome in *M. abscessus* has not yet been sufficiently studied, and it is currently unclear whether synergy correlates with positive clinical outcomes (71). This challenge in linking *in vitro* measurement to *in vivo* response is not unique to NTMs and is well characterized in tuberculosis (72, 73). However, previous studies have shown that measurement of the drug combination response across multiple growth conditions that resemble the host’s microenvironment is predictive of treatment shortening in Mtb (30, 74). Environmental nutrients have a critical effect on drug interaction in both *M. abscessus* and Mtb, and antibiotic response is altered when grown in artificial cystic fibrosis sputum (30, 75, 76). We, therefore, expect that making drug interaction measurements in different host-mimicking growth conditions and testing which of these growth environments predicts treatment outcomes in preclinical animal models and in humans is a critical step in drug combination design for *M. abscessus* and other NTMs. Together, the dependence of drug interactions on strain and variation in drug response with strain and growth conditions suggests that we have failed to identify a universal combination therapy that is effective for *M. abscessus* not because good drug combinations do not exist, but rather because we need to better understand how to tailor combination therapy to each infection.

## MATERIALS AND METHODS

### Antimicrobials

The antimicrobials agents used in this study (except for SPR719) were obtained from Sigma Aldrich. SPR719 was a gift from Spero Therapeutics. Stock solutions were prepared in DMSO or sterile water + 0.01% Triton X-100, depending on solubility, and stored in single-use aliquots at −20°C until used.

### Strain and culturing

Measurement was made using *M. abscessus* subsp. *abscessus* strain ATCC19977 (reference strain) and clinical isolates obtained from patient sputum at Tufts Medical Center, Infectious Disease Clinic (TMC1, TMC2, and TMC3) (77). All strains were cultured in a 7H9 medium supplemented with 0.05% Tween 80, 0.2% Glycerol, and 10% BBL Middlebrook ADC enrichment. Cultures were started from frozen aliquots and allowed to grow to the mid-log phase (OD_600_ between 0.4 - 0.6), shaking at 37°C overnight. Cultures were then diluted once to the lag phase (OD_600_ between 0.05 - 0.1) and allowed to grow to the mid-log phase before performing assays.

### DiaMOND measurement

DiaMOND was used to measure drug interactions, as previously described for *Mycobacterium tuberculosis* (18). Details of DiaMOND can be obtained in Mycobacteria Protocol 4^th^, chapter 30^th^ (78). Briefly, DiaMOND uses equipotent drug-combination dose-response curves to approximate the shape of checkerboard isoboles at the same level of growth inhibition. Minor adjustments (e.g., the dose-response curve is centered around 50% inhibitory concentration (IC_50_) in contrast to 90% inhibitory concentration (IC_90_) for *M. tuberculosis*, the modification of fractional inhibition concentration to fold shifts in inhibition concentration) were made to account for the rapid growth of *M. abscessus* and low drug potency relative to *M. tuberculosis*. Details of these adjustments are explained in subsequent sections.

#### Growth inhibition assay

Antibiotics tested were either dissolved in dimethyl sulfoxide (DMSO) or sterile water + 0.01% Triton X-100 and stored at −20°C in single-use aliquots. Assays were performed in clear, flat bottom 384-well microplates. Drugs were dispensed using a digital drug dispenser (D300e Digital Dispenser, HP). Drug wells were randomized across plates to minimize plate position effects. Bacterial cultures were diluted to OD_600_ of 0.05 in fresh medium, and 50μL of diluted culture was added to each well for drug treatment. Plates were sealed with optically clear plate seals and incubated without shaking at 37°C. Growth (OD_600_) was measured by a microplate reader (BioTek) 48h after drug treatment.

#### Data analysis

OD_600_ data were processed using MATLAB’s custom analysis pipeline (MathWorks). Data were first derandomized from the plate layout, and dose-response curves were organized for each drug and drug combination. The first row of wells in each 384-well plate contained medium-only wells. The median of these medium-only wells was subtracted from each well in a plate as a background. Drug-treated wells were normalized to the mean of the untreated wells (controls), and the values obtained were subtracted from 1 to get a dose-response inhibition curve where 0 and 1 represented no growth inhibition and full growth inhibition, respectively. Growth inhibition curves were then fit to a three-parameter Hill curve using a nonlinear solver in MATLAB (79). The fit accuracy was assessed using the R^2^ metric. Equations derived from the Hill function were used to calculate different inhibitory concentration values (IC values) along the dose-response curve.

#### Drug interaction calculation

To assess whether drugs in combination were synergistic, additive, or antagonistic, we measured the fractional inhibitory concentrations (FIC). We used Loewe additivity as the null model to calculate the FIC values (80). We calculated the expected IC value of the combination AB (drug A with drug B) from the intersection of the combination line to the line of additivity determined by the IC values of A and B alone (Fig. 1A). Finally, the FIC value is calculated by dividing the experimentally observed IC by the expected IC value of the drug combination.

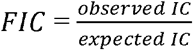

We report the log_2_ of the FIC so that the magnitude of synergy and antagonism scores are balanced around zero. A log_2_FIC value < 0 is synergistic, log_2_FIC value ~= 0 is additive, and log_2_FIC value > 0 is antagonistic. FIC scores were calculated at IC_50_ (FIC_50_) and IC_90_ (FIC_90_). With DiaMOND, we can measure drug interaction scores using other null models such as Bliss independence. By Bliss independence, the expected IC value for [AB] is calculated by multiplying drug A’s effect and drug B’s effect at the desired IC level.

Certain drugs were tested at a fixed concentration (instead of increasing doses) due to their lack of inhibitory effect as a single agent. These drugs are also referred to as potentiator candidates. The concentration of the potentiator candidates was estimated from previously reported serum concentrations. To quantify drug interaction scores for potentiators, fold shifts in ICs (FsICs) were calculated as a ratio of the observed IC_50_ value of a drug pair combination to the IC_50_ of the active, non-potentiating drug (Fig. 1B). Log_2_FsIC < 0 are potentiated whereas log_2_FsIC > 0 are attenuated.

#### Quality control

Assay quality was assessed in three different ways. The Z-factor was used to determine the quality of the assay at the plate level. For a given plate, the Z-factor was calculated as:

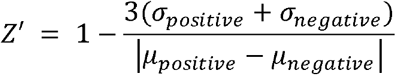

where *σ* is the standard deviation of the positive and negative control (untreated cells), *μ* is the mean of the positive and negative controls. Plates with a *Z′* between 0.5 - 1 indicate excellent assay with a statistically reliable separation between positive and negative control.

The second level of assessment was at the dose-response level. The R^2^ value derived from fits of the dose-response curves to Hill curves was used to determine fit accuracy. We combined R^2^ values with a visual inspection of the fits. Any Hill curve fits with R^2^ values below 0.7 were marked as poor fits and rejected from further analysis. Because of the intrinsic resistance property of *M. abscessus*, it is challenging to obtain consistent dose-response curves (81). For example, to capture data points close enough to each other to draw an accurate dose response, we need to design doses to increase by 1.5x instead of 2c, which limits the testing range. In addition, the noise of the dose-response curve made fitting challenging. Finally, obtaining a maximum inhibitory concentration (MIC) for all drugs is difficult due to the heterogeneity of *M. abscessus* (81). To overcome these difficulties, we designed the experiment around IC_50_ (instead of IC_90_), which is achieved for most drugs and is more reproducible than IC_90_ (Table S1). To assess whether doses were sampled in an equipotent manner for combination dose-response, the angle of the combination dose-response was calculated (the ideal angle is 45°). Angle deviation beyond 22.5° of this ideal equipotent dose-response was deemed too far for the approximation of isoboles in the checkerboard and was eliminated from the analysis.

### Clinical isolate growth rate measurement

To measure the growth rate of clinical isolates obtained from patient sputum at the Tufts Medical Center Infectious Disease Clinic (TMC1, TMC2, and TMC3), suspended culture of these isolates were grown in a 7H9 medium supplemented with 0.05% Tween 80, 0.2% Glycerol, and 10% BBL Middlebrook ADC enrichment. Cultures were started from frozen aliquots and allowed to grow to the mid-log phase (OD600 between 0.4 - 0.6), shaking at 37°C overnight. Cultures were then diluted once to the lag phase and allowed to grow to the mid-log phase again before performing the assay. Growth rate measurement was performed in a 96-well microplate, with five biological replicates per isolate. Reference strain ATCC19977 was also included for reference purposes. 150μL of culture at OD_600_ around 0.05 was dispensed into each well. The microplate was sealed with an optically clear plate seal and incubated inside a plate reader at 37°C. OD_600_ was recorded for 18 hours in 30 minutes intervals. Doubling time was calculated using OD_600_ values closest to the log phase. Doubling time was calculated as:

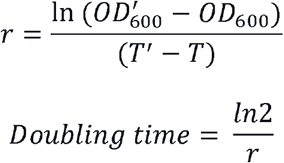

## ACKNOWLEDGEMENTS

We thank members of the Aldridge laboratory, Veronique Dartois, Alyssa Greig, and Aisling Lavelle for insightful discussion. This work was supported in part by the Stuart B. Levy Center for Integrated Management of Antimicrobial Resistance at Tufts (Levy CIMAR), a collaboration of Tufts Medical Center and the Tufts University Office of the Vice Provost for Research Research and Scholarship Strategic Plan. BBA was supported, in part, by NIH 5R01AI150684 (to BBA). FPM was supported by a financial grant from Mukoviszidose Institut gGmbH, Bonn, the research and development arm of the German Cystic Fibrosis Association Mukoviszidose e.V (project number 2004 – FM). JK was supported by National Institution of General Medical Science (T32GM008448). PAL was supported by National Institution of General Medical Science (T32AI007422).

## AUTHOR CONTRIBUTION

NV, YD, JLF, TS, and BBA conceived and designed the experiments. NV, YD, JK, and PL performed the experiments. NV, YD, and BBA conceived and designed the computational analysis. NV and YD performed the computational analysis. The manuscript was written by NV, YD, TS, and BBA. All authors contributed to the technical interpretation, interpretation of the results, and the editing of the manuscript.

