## Supplemental Figures for "Novel synergies and isolate specificities in the drug interactions landscape of *Mycobacterium abscessus*"

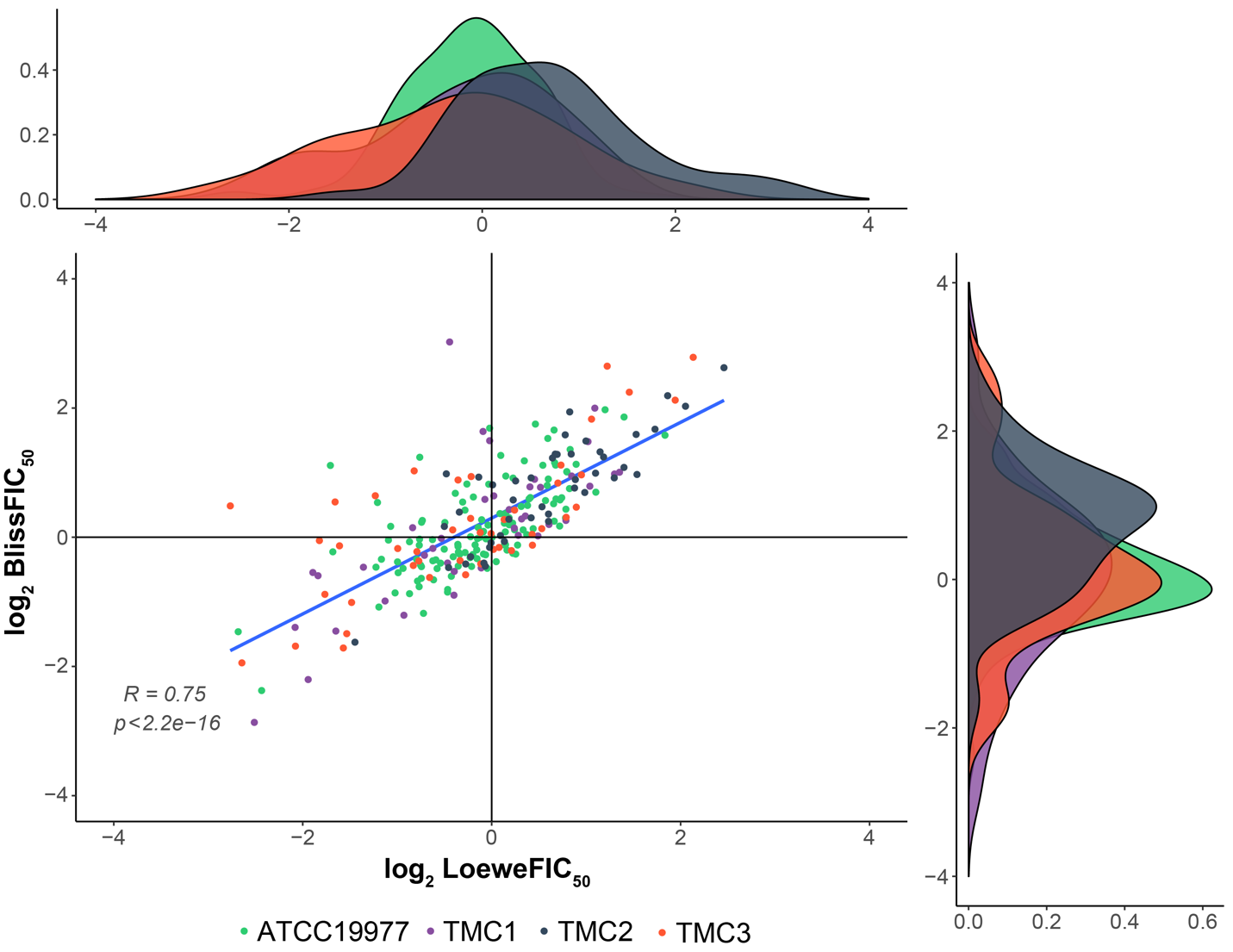

**FIG S1. Correlation between LoeweFIC_50_ and BlissFIC_50_**. Scatter plot represents the correlation between log_2_FIC_50_ calculated using Bliss independent (y-axis) versus log_2_FIC_50_ calculated Loewe additivity (x-axis). Pearson correlation coefficient R and p-value for the corresponding test of correlation are reported. The linear regression line is provided for visualization. The distribution of log_2_FIC_50_ values calculated using the two methods mentioned above is represented by the plot on the top (log_2_FIC_50_ calculated Loewe additivity) and the plot on the right (log_2_FIC_50_ calculated using Bliss independent). The density of the distribution is on the y-axis, and the log_2_FIC_50_ value is on the x-axis. Different colors in both plots indicate different strains used in the study.

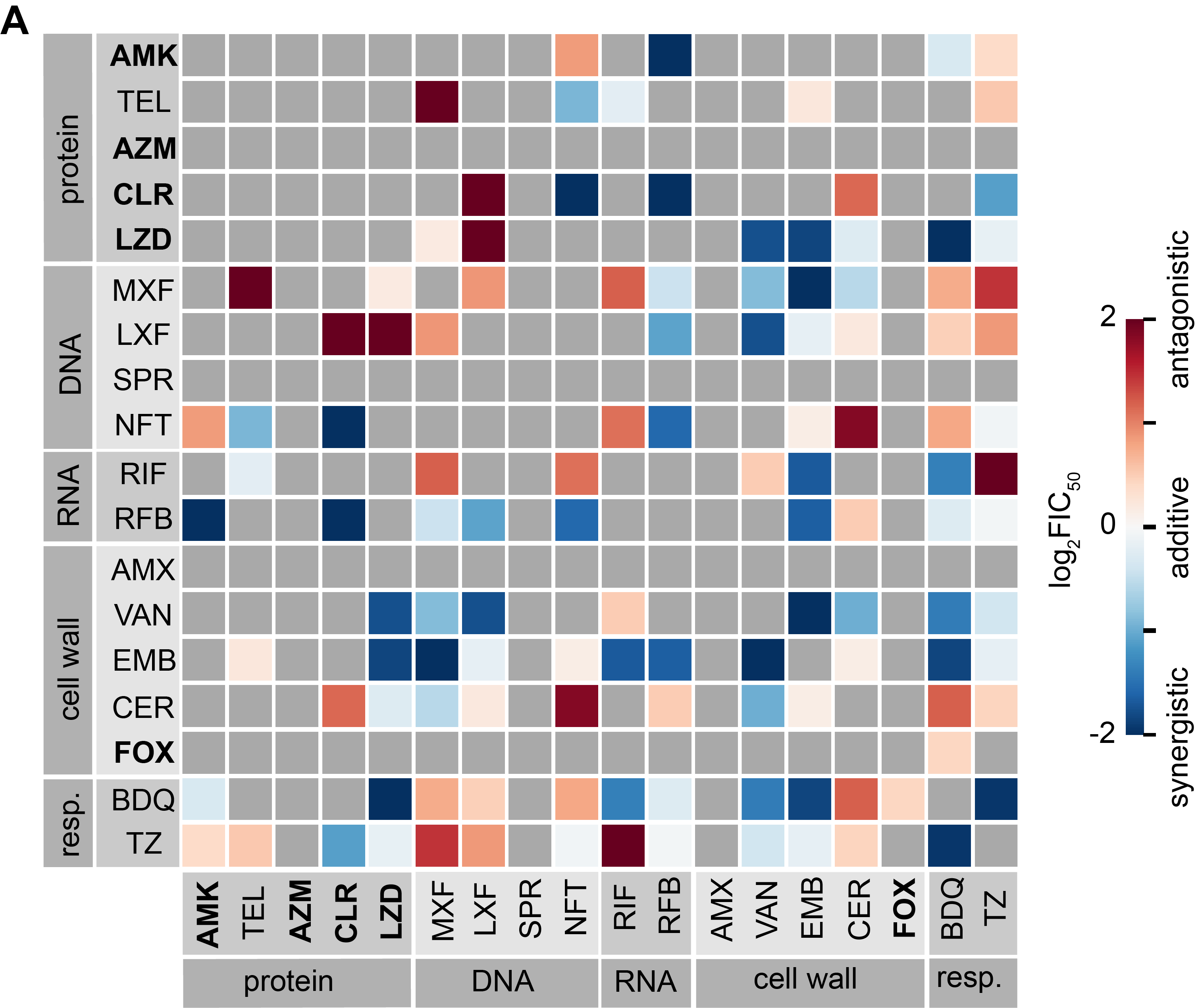

**FIG S2. Drug interaction landscape of *M. abscessus* strain ATCC19977 at IC_90_**. Heatmap of pairwise drug interactions among 18 drugs at IC_90_. Drugs are organized by the mechanism of action, and drugs recommended for the treatment of *M. abscessus* infection are indicated in bold text. Drug interactions are evaluated with log_2_FIC_90_ values: log_2_FIC < 0 (synergy, blue) and log_2_FIC > 0 (antagonism, red). Gray boxes indicate unmeasurable drug combinations due to poor reproducibility.

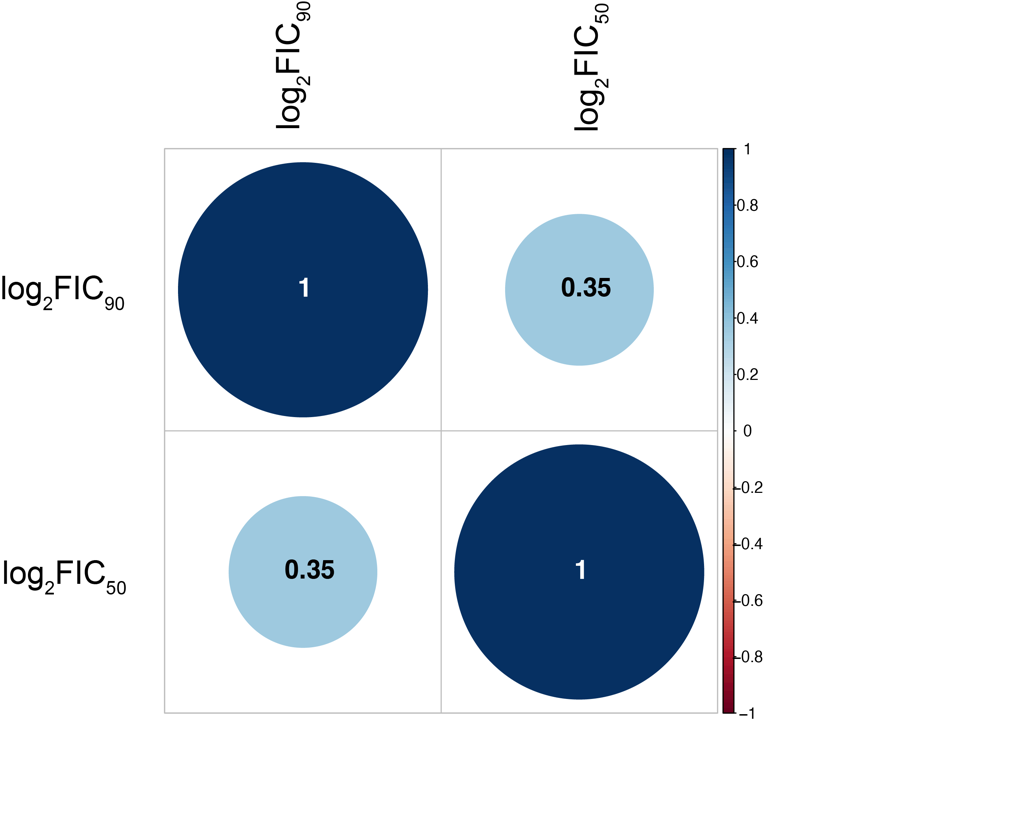

**FIG S3. Correlation between log_2_FIC_50_ and log_2_FIC_90_ in ATCC19977**. Correlation plot represents the correlation between log_2_FIC_50_ and log_2_FIC_90_ in ATCC19977. Pearson correlation coefficient R for the corresponding test of correlation is reported by the number inside the bubble. The color of the bubble also indicates the strength of correlation, with blue indicating positive correlation (maximum value of 1) and red indicating negative correlation (maximum value of -1).

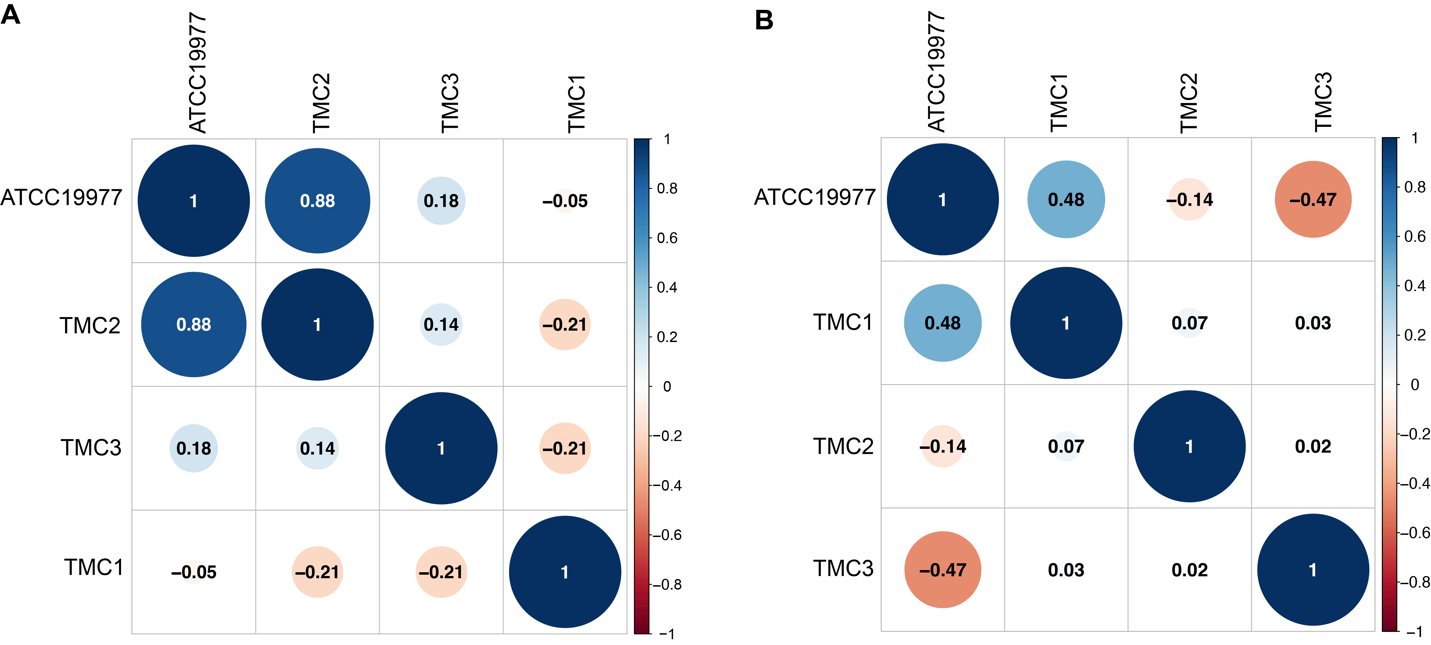

**FIG S4. Correlation of IC_50_ and log_2_FIC_50_ between strains used in the study**. Correlation plot represents the correlation of IC_50_ (A) and log_2_FIC_50_ (B) between lab strains ATCC19977, TMC1, TMC2, and TMC3. Pearson correlation coefficient R for the corresponding test of correlation is reported by the number inside the bubble. The color of the bubble also indicates the strength of correlation, with blue indicating positive correlation (maximum value of 1) and red indicating negative correlation (maximum value of -1).
